# Spatial mapping of acoustic impedance of shrimp scale using multiple overlap discrete wavelet transform (moDWT)

**DOI:** 10.1101/2023.10.14.562137

**Authors:** Shivam Ojha, Komal Agarwal, M Sarim Ameed Khan, Amit Shelke, Anowarul Habib

**Author notes:** these authors have contributed equally to this work and are co-first authors (Shivam Ojha, Komal Agarwal).

## Abstract

There are lots of challenges associated with conventional optical observation of biological tissues, where specimens are typically sliced and stained for better contrast. In contrast, Scanning Acoustic Microscopy (SAM) is a versatile label-free imaging technology widely applied in various domains, including biomedical imaging, non-destructive testing, and material research. It excels in offering precise visualization of both surface and subsurface structures, providing valuable insights through visual inspection and quantitative analysis. Acknowledging the SAM, this paper presents acoustic impedance microscopy of the shrimp scale in a novel manner. The proposed technique aims to image the local distribution of cross-sectional acoustic impedance in biological tissue, which is a parameter closely related to sound speed and potentially valuable for tissue characterization. The study leverages advanced signal processing techniques, maximal overlap discrete wavelet transform (moDWT), to decompose acoustic responses effectively. The moDWT, with its ability to handle signals of various lengths without constraints, is highlighted as a promising approach. To determine shrimp scale impedance, we first establish the accuracy of the proposed algorithm using PVDF as the target and polyimide as reference material. The results indicate an algorithm accuracy exceeding 90%. An impedance map is generated through Gaussian process regression (GPR), which predicts the impedance over the complete domain, addressing spatial variations in biological specimens. The resulting acoustic impedance maps provide in-depth insights into the functional framework and advance our understanding of shrimp biomechanics.

## 1 Introduction

Scanning Acoustic Microscopy (SAM) is a valuable label-free imaging technique utilized in diverse fields such as biomedical imaging, non-destructive testing, and material research^1–5^. It enables the precise visualization of both surface and sub-surface structures. SAM offers both visual inspection and precise quantitative analysis capabilities, making it essential in material research, non-destructive testing, and biomedical imaging. Its non-invasive nature facilitates detailed micro-structural characterization and assessment of mechanical properties, defect detection, and monitoring of the health of composite structures^6–8^.

SAM’s versatility positions it as a vital tool for applications requiring accurate imaging and analysis. The concept of biomimicry, drawing inspiration from nature’s diverse materials, has gained traction. Materials like bones, teeth, and nacre inspire innovative engineering solutions. Researchers are exploring biomimicry in materials with lower mineral content, such as fish scales, crustacean exoskeletons, and osteoderms, blending flexibility, strength, and toughness. Chitin, the second most abundant natural polymer, plays a crucial role in the exoskeletons of organisms like shrimps, providing insights into structural impacts on functionality^9^.

Conventional optical observation of biological tissues faces challenges, with SAM offering a non-destructive alternative for rapid observations. SAM utilizes focused acoustic signals, emerging as a powerful tool for tissue characterization and elastic parameter imaging^10–13^. This paper introduces acoustic impedance microscopy of shrimp scales as a non-invasive method, aiming to image local acoustic impedance distribution related to sound speed, which is crucial for tissue characterization. The proposed technique aims to image the local distribution of cross-sectional acoustic impedance in biological tissue, which is a parameter closely related to sound speed and potentially valuable for tissue characterization. By exploiting the relationship between acoustic impedance, sound speed, and density, this methodology enables micro-scale imaging through acoustic response.

The acoustic response contains a bunch of frequencies that evolve with time, giving rise to the need for decomposition using time-frequency analysis. The most basic methods for time-frequency analysis are time windowing and Fourier analysis, which focus on breaking down signals into respective temporal and frequency components^14^. Both of these methods face difficulty in examining the signal components presented in multiple time scales. Time windowing requires careful selection of appropriate averaging time, while Fourier analysis involves pre-processing steps like data windowing. Advanced tools like short-term Fourier transform^15^ (STFT) and Wavelet transform have better signal decomposition in the multi-scale resolution^14^. Within the domain of wavelet decomposition, Discrete Wavelet Transform (DWT) excels in decomposing signals into both time and frequency domains^16,17^. However, the DWT imposes restrictions on the length of signals, which should be multiple powers of two. This limitation restricts the DWT application, and decomposition depends on whether the event span falls within a wavelet averaging window or not. The more advanced version of DWT, the maximal overlap discrete wavelet transform (moDWT), retains down-sampled values at each decomposition level and does not put restriction on signal length^14,17^. Considering the benefits of the moDWT over other methods, it is suggested that combining the moDWT with soft computing models can offer a more effective and efficient approach for extracting the characteristic features through the acoustic response of shrimp scales. The article presents the process of extracting dominant frequencies from the acoustic signal through moDWT combined with bandpass filter, thus building up a basis for the estimation of impedance. Further, Gaussian process regression (GPR) - based kriging is employed for spatial interpolation of impedance. The GPR is very flexible and can incorporate the uncertainty in the observed data while performing regression and handles complexities in non-linear mapping^18^. The algorithm is validated on known materials before application to bio-samples. Figure 1 depicts the overall strategy used in this paper.

**Figure 1.**
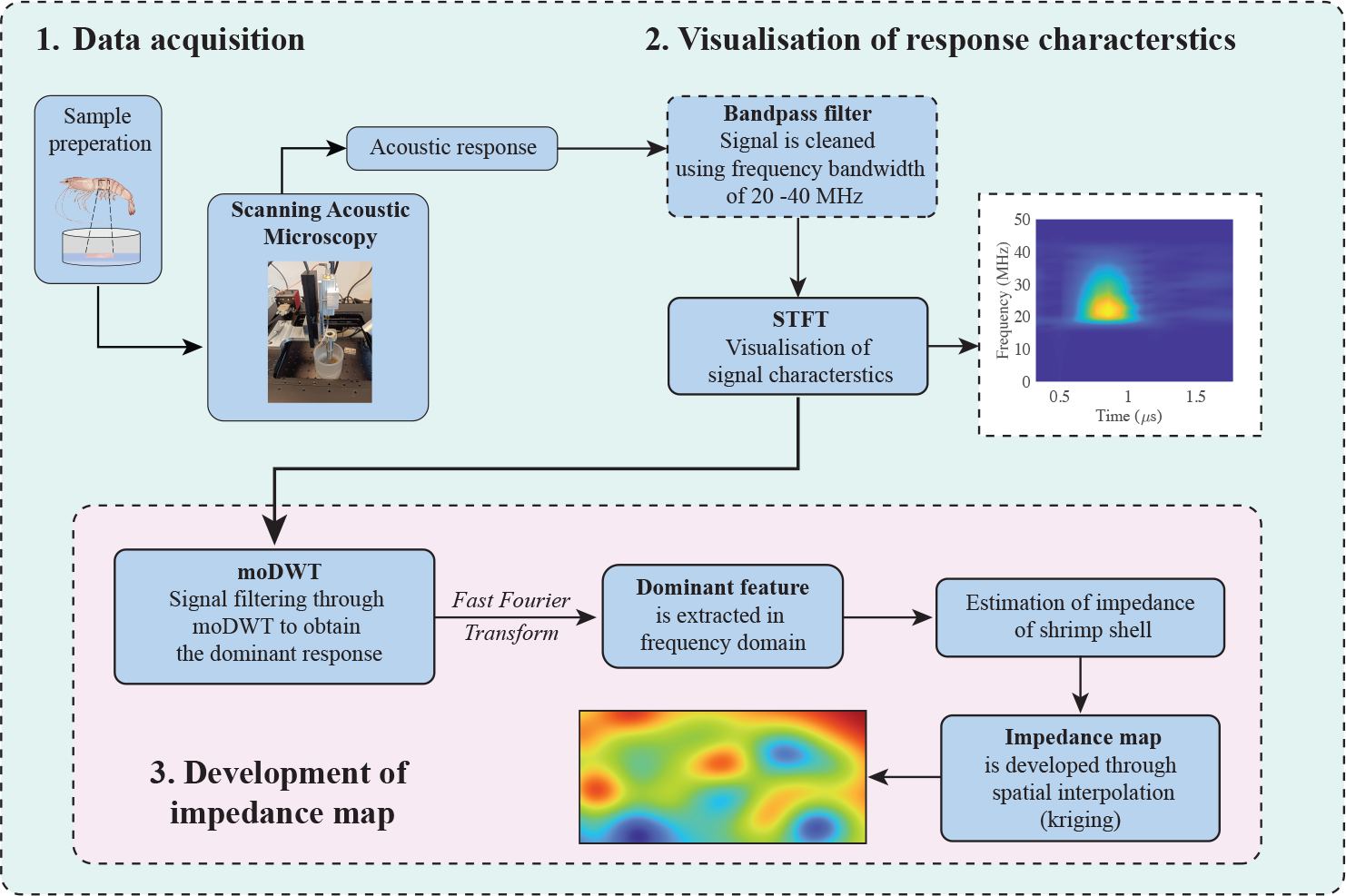
The figure illustrates the complete strategy for the development of the impedance map of the shrimp scale. The first section involves SAM imaging, where Scanning Acoustic Microscopy is used to capture images of several samples. In the second section, we visualize and analyze the acoustic response characteristics of the samples. The third and final section is dedicated to impedance mapping, which entails mapping and calculating impedance values based on the acquired data.

This study meticulously explores the biomechanical attributes of shrimp using high-resolution scanning acoustic microscopy. The non-invasive methodology delves into the intricate biomechanical properties of the shrimp’s scale, laying the foundation for future research. The objective is to establish innovative non-invasive examination techniques for early detection of structural alterations within the shrimp community. The study systematically experiments with high-frequency acoustic waves penetrating shrimp scales, determining acoustic impedance through the proposed algorithm. Visual representations in the form of acoustic impedance maps effectively reveal the functional utility and intricate biomechanical framework of distinct components within the shrimp’s exoskeleton. This pioneering approach aims to deepen our understanding of shrimp biomechanics, offering improved insights into the broader marine ecosystem and revolutionizing the diagnosis of structural changes within marine communities.

## 2 Experimental procedure

### 2.1 Sample preparation

The shrimp scales were obtained from healthy shrimps from the arctic water collected by the Department of Fisheries at UiT, Norway. The shrimp scales were plucked fresh and carefully from the edge of the scale using broad tweezers to avoid any cracking.

A solution consisting of 2 wt% Agarose (ultra-low gelling temperature, Sigma Aldrich) was prepared by dissolving it in 10 ml of distilled water. The mixture was agitated in a beaker using a magnetic stirrer at a temperature of 100 °C for a duration of 10 minutes. Subsequently, a thick gel was poured onto a standard Petri dish (μ-Dish 35 mm from Ibidi), and atop the gel, a shrimp scale was positioned and gently pressed down into the agarose layer. A visual representation of this sample preparation process can be observed in Figure 2 (a and b). Cold distilled water was carefully added to the Petri dish, and the imaging procedure using SAM was promptly initiated to ensure the preservation of the sample’s integrity. The inclusion of agarose served the dual purpose of maintaining the sample’s freshness and securing the scale in position within the water bath^19^. Furthermore, agarose exhibits an acoustic impedance similar to that of water^20^. To measure the acoustic impedance of the shrimp scale, two other known sample impedances polyimide (PI) and polyvinylidene fluoride (PVDF) were also embedded in the agarose gel along with the shrimp scale as shown in Figure 2c. This approach allowed us to capture images of all the samples together in one session, facilitating a comparative analysis.

**Figure 2.**
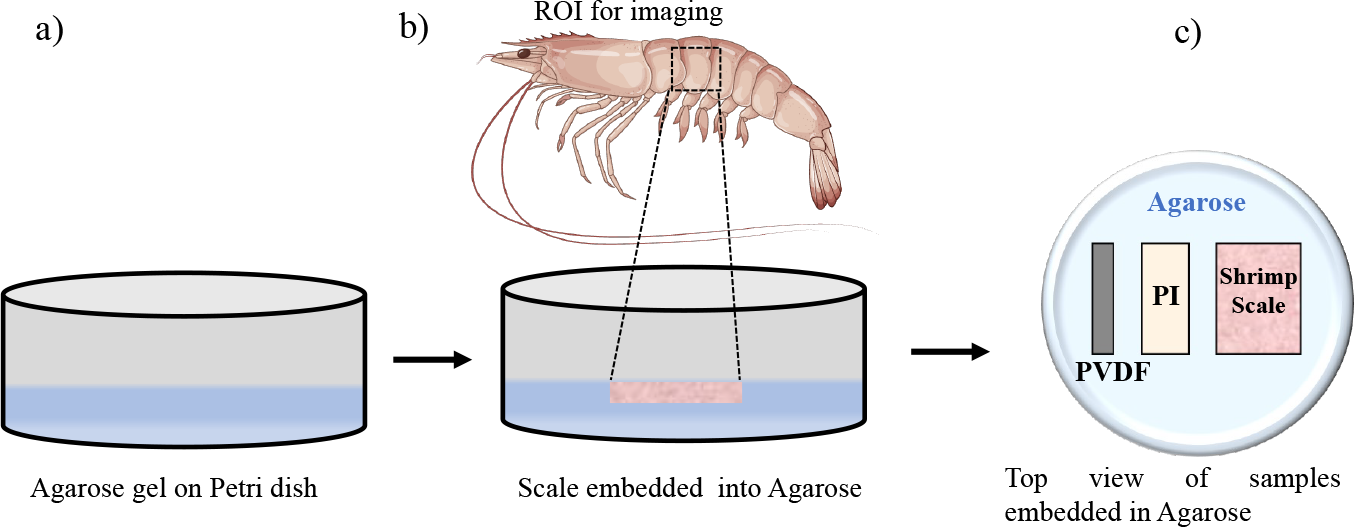
A schematic representation of sample preparation provides a diagrammatic depiction of the process and steps involved in preparing a sample for experimentation. Figure (a) represents a thick Agarose gel layer into a regular Petri dish, while in Figure (b), a shrimp scale is carefully placed and pressed into the Agarose layer. The area of interest (ROI) was then imaged using SAM. Figure (c) depicts a schematic representation featuring the reference samples polyvinylidene fluoride (PVDF) and Polyimide (PI), as well as the target sample shrimp scale.

### 2.2 Scanning acoustic microscopic imaging

Figure 3 demonstrates a labeled representation of a SAM, employed for acquiring images of samples. SAM employs both reflection and transmission modes to generate images, each offering unique insights into different aspects of the sample’s characteristics. The image of SAM, as depicted in Figure 2, has been annotated to highlight its key components or operational settings and serves as a reference for image acquisition. More comprehensive explanations of the operational principles for these modes can be found in the following literature^21,22^.

**Figure 3.**
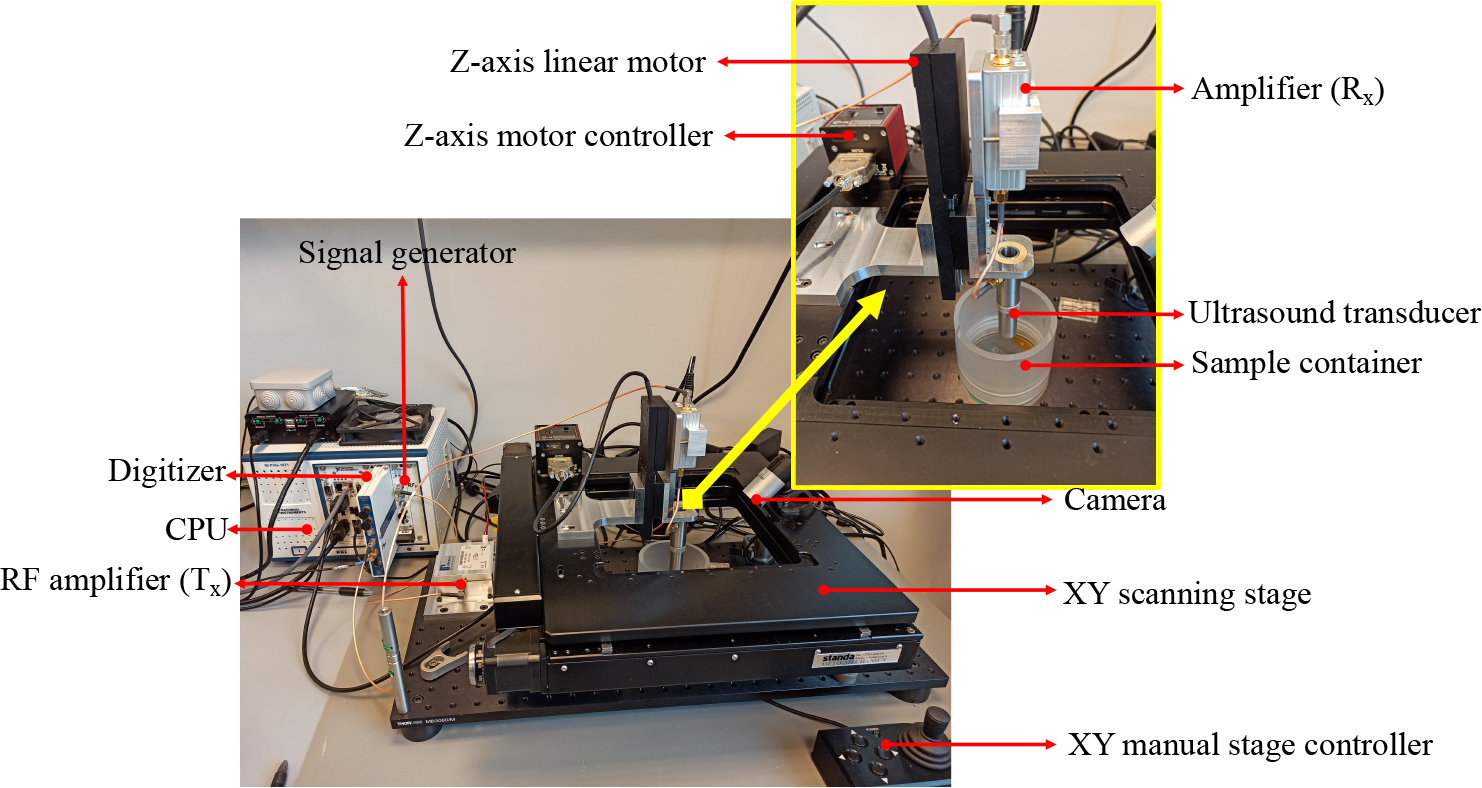
Figure depicts a labeled image of SAM used for image acquisition as discussed in this paper. The experimental setup demonstrates all the fundamental components that constitute a SAM.

In this article, our emphasis lies in the utilization of the reflection mode for scanning samples. To achieve this, a commonly employed technique involves using a concave spherical sapphire lens rod to focus acoustic energy through a coupling medium, typically water. Ultrasonic signals, generated from a signal generator, are directed toward the sample. Upon reflection from the sample’s surface, the reflected waves are detected and transformed into a digital signal, commonly referred to as an A-scan or amplitude scan.

To generate a C-scan image of the sample, this process is repeated at various positions within the XY plane. Another perspective on visualizing a C-scan is to regard it as the integration of A-scans in two dimensions. A custom-built SAM, as illustrated in Figure 2, was controlled by a LabVIEW program. This SAM system featured a Standa high-precision scanning stage (8MTF-200-Motorized XY Microscope Stage) for gathering experimental data. Acoustic imaging capabilities were enabled using National Instruments’ PXIe FPGA modules and FlexRIO hardware housed in a PXIe chassis (PXIe-1082) with an integrated arbitrary waveform generator (AT-1212)^21–23^. The transducer was excited with Mexican hat signals and amplified using an RF amplifier (AMP018032-T). Acoustic reflections from the sample were then amplified, and digitized at a high speed (1.6 GS/s) with a 12-bit digitizer (NI-5772). A 30 MHz PVDF-focused Olympus transducer with a 6.35 mm aperture and 12 mm focal length was employed^24^. The thickness of the films was measured using a digital micrometer. The PVDF, PI film, and the shrimp scale were approximately 105, 130, and 100 *μm* thick, respectively. The scan area (x, y) of 10 mm *×* 2 mm was imaged with an x, y pixel size of 50 *μm*. The samples were scanned from above 200 *μm* till 800 *μm* below the focal point of the thickest sample (PI), with a step size of 50 *μm*. This was to ensure that all the samples with varying thicknesses were imaged in their focal plane and that the z-scans of the samples were obtained at the different planes.

## 3 Theoretical background

### 3.1 Mechanics of the acoustic waves

Acoustic microscopy operates non-destructively and penetrates acoustic waves to make images of internal features visible. This propagation of acoustic waves generates the normal and shear stress in the medium, which plays a fundamental role in the classification of acoustic waves into P waves (Primary or Pressure waves) and S waves (Secondary or Shear waves). For

the planar interface, the incident of the P-wave gives rise to four distinct wave components; a) reflected P wave (RPP), b) reflected S wave (RPS), c) transmitted P wave (TPP), and d) transmitted S wave (TPS). To describe these wave phenomena mathematically, the wave potentials can be written as follows^25^:

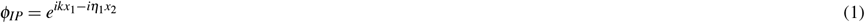

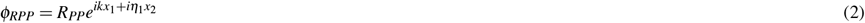

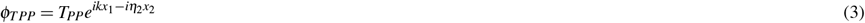

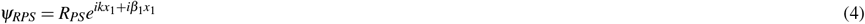

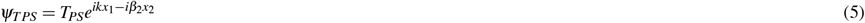

where subscript numbers 1 and 2 represent the medium and,

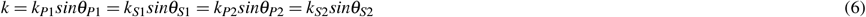

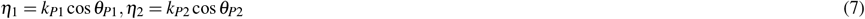

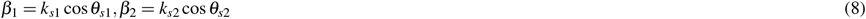

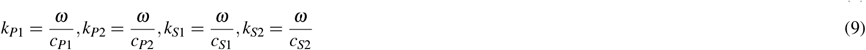

where, *k*_*P*1_, *k*_*P*2_, *k*_*S*1_, *k*_*S*2_ are the wave numbers of P and S waves traveling in material 1 and 2, and other symbols have their standard meanings. Considering the special case of normal incidence of P-wave, i.e., *θ*_*P*1_ = 0, *θ*_*P*2_ = 0, *θ*_*S*1_ = 0, *θ*_*S*2_ = 0, through the continuity of the displacements and the stress conditions at the interface, the following simplified matrix is obtained as follows:

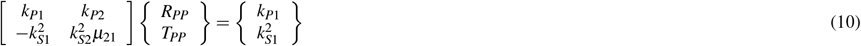

Here, *μ*_21_ = *μ*_2_*/μ*_1_ is the ratio of lame’s constants, and the values of reflectance *R*_*PP*_ are obtained as follows:

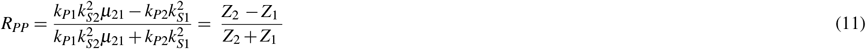

where, *Z*_1_ and *Z*_2_ are the acoustic impedance of the medium 1 and 2, respectively.

### Estimation of Impedance

Let the characterization frequency of the transmitted/incident signal be *S*_0_, then the corresponding frequency of reflected response of reference and target medium is represented by *S*_*re f*_ and *S*_*target*_. Using the property of reflectance, the relation between the characterization of the transmitted signal and the reflected response of the target medium is written as^25–27^:

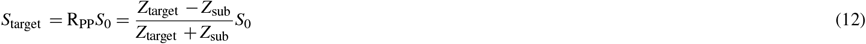

where *Z*_*target*_ and *Z*_*sub*_ are the mean value of the acoustic impedance of the target and substrate, respectively. Similarly, the same can be written for the reflected response of the reference medium as

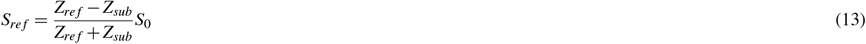

where *Z*_*re f*_ is the mean value of the acoustic impedance of the reference medium. The measurement is possible only for the *S*_*re f*_ and *S*_*target*_ and *S*_0_ cannot be measured directly. The acoustic impedance of the target medium is subsequently calculated as a solution using Eq. 12 and Eq. 13.

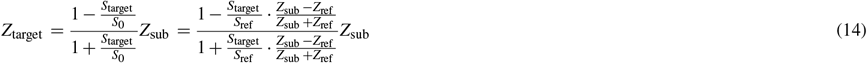

However, the identification of ratio *S*_*target*_*/S*_*re f*_ is quite challenging, and therefore, measurement error is always present in such cases. Thus, the current work considers estimated impedance as noisy observations while performing spatial interpolation to develop the impedance map over the complete domain.

### 3.2 Discrete wavelet transform (DWT)

DWT is a time-frequency analysis technique that decomposes the original signal into approximation and detailed coefficients through filtering and downsampling. The signal is decomposed into multiple levels, and each level is characterized through its approximation and detailed coefficients. This section just outlines the fundamental concept of DWT. Interested readers can refer to the following literature^14,16,17^.

The practical implementation of DWT is carried out using Mallat’s algorithm which uses a pair of filters: a low-pass filter (scaling function) and a high-pass filter (wavelet function). The process is then recursively applied to the resulting low-pass signal. This recursive application on low-pass signals creates different levels of wavelet decomposition. For a discrete signal, **X** = *{X*_*t*_, *t* = 0, 1, *…*, *N −* 1*}*, the DWT computes the wavelet coefficient for the discrete wavelet of scale 2 ^*j*^ and location 2 ^*j*^*k* using the following equation:

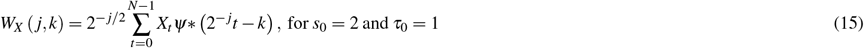

where *W*_*X*_ (*j, k*) is the wavelet coefficient and *N* is an integer power of two. In order to reconstruct the filtered signal, the necessary levels are selected. The wavelet coefficients associated with those levels were passed through inverse discrete wavelet to obtain the filtered response. Further, DWT lacks shift-invariance and has limited frequency resolution in higher scales due to down-sampling. The improved variant called maximal overlap discrete wavelet transform (moDWT) is more suitable for time-frequency analysis where precise localization is crucial. The current work utilizes the moDWT for characterizing the signal which is described in the subsequent section.

### Maximal overlap discrete wavelet transform (moDWT)

It divides the frequency band of the input signal into scaling and wavelet coefficients using low- and high-pass filters, that is, scaling and wavelet filters. moDWT can be properly defined for arbitrary signal length, while the DWT is limited to a signal length with an integer multiple of a power of two^14^. Further, moDWT achieves redundancy through an oversampled representation which enables more accurate statistical analysis. Details on moDWT can be found in Percival and Walden^17^.

Let *X* = *X*_*t*_, *t* = 0, …, *T −* 1, be the time series data, then, the j^th^ level wavelet and scaling filters are denoted as 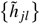 and 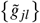, respectively. The scaling and wavelet coefficients calculated by the Mallat algorithm are described as follows:

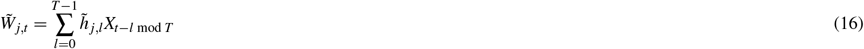

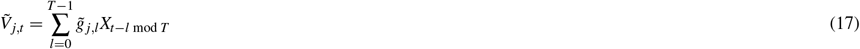

where *j* = 1, 2, …, *J*_0_ is the level of wavelet decomposition, and ‘mod’ denotes the remainder of dividing two numbers. The wavelet 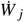 and scaling 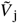 coefficients vectors of moDWT are written as:

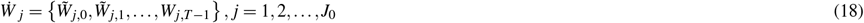

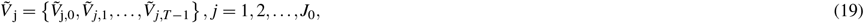

where 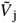 and 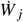 are related to the smallest and highest frequency components of the original signal. 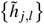, and 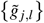 are the j^th^ level moDWT high-pass filter 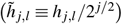 and low-pass filter 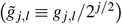 and *J*_0_ is the highest decomposition level. The filters are determined depending on the mother wavelets. For the level 3 decomposition, the moDWT decomposes an original signal *X* into a low-pass filtered approximation component (A_3_) and high-pass filtered detail components (*D*_1_, *D*_2_ and *D*_3_). The equations of moDWT-based multi-resolution analysis can be used for the reconstruction of decomposed signal and are written as follows:

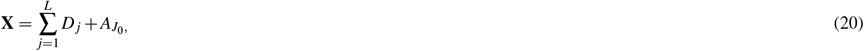

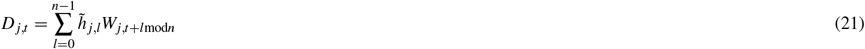

### 3.3 Spatial interpolation through GPR-based kriging

Kriging^28^, also known as Gaussian process regression (GPR), is a method of interpolation based on the Gaussian process. It develops a meta-model of a partially observed function (of time and/or space) with an assumption that this function is a realization of a Gaussian process (GP)^18,29^. In GPR, the prescribed forms of mean and covariance functions are assumed, and the hyperparameters of the kernel functions are computed from observation via maximizing the log-marginal likelihood function.

For kriging, the observations are modeled as a GP with constant mean and prescribed kernel function. Let the known locations be represented by 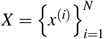 where *x* is the two-dimensional vector and the observed state at these locations isgiven by *y* = {*y*^(1)^, *y*^(2)^, …, *y*^(*N*)^}. Let the meta-model between the observed state and known locations is given by:

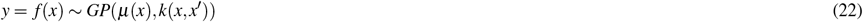

where *μ*(*x*) is the mean function and *k*(*x, x*′) is the kernel or covariance function. By definition, the prior of the observed state vector is Gaussian and is given by *p* (*f* (*x*) | *x*) = *N* (*f* (*x*) | *μ, K*). Given the prediction locations as *x*_*∗*_, the predicted state *y*_*∗*_ = *f* (*x*_*∗*_) is also jointly gaussian and is given by

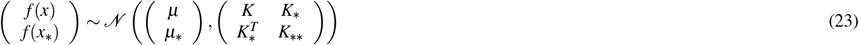

Using Bayesian transformation, the posterior mean and variance of the predicted state are written as

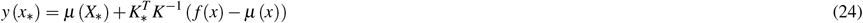

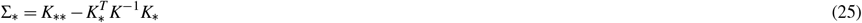

## 4 Proposed algorithm

In the current work, the impedance map of the shrimp scale is developed through acoustic microscopy. The reflected response of the acoustic imaging contains multiple waves having different frequency components that are transmitted and reflected through the multiple interfaces resulting in the evolution of multiple frequencies with time. The reflected signal is first analyzed through the Short-Time Fourier Transform (STFT), which computes the Fourier transform of small, overlapping time segments of a signal, revealing the signal’s frequency content as a function of time^15^. The article utilizes the STFT for observing the frequency variations and presents the following algorithm for the extraction of the desired frequency content and the development of an impedance map over the complete domain.

**Step 1: Removal of noise through band-pass filter**

The incident P wave used in the experiment is the ricker wave with a dominant frequency of 30 MHz. The reflected wave obtained contains unwanted frequencies arising from noise which need to be removed first. This cleaning of the signal is achieved with a bandpass filter.

**Step 2: Filtering of cleaned signal through moDWT**

Utilizing the merits of the moDWT, the cleaned signal is decomposed into a series of wavelet and scaling coefficients with each level. The filtered signal is reconstructed through inverse moDWT using the essential wavelet and scaling coefficients

**Step 3: Selection of dominant frequency**

The filtered signal that has been reconstructed through moDWT is transformed into a frequency domain through the fast Fourier transform. In the frequency domain, the frequency corresponding to the maximum amplitude is assumed to be the signal-characterizing feature and is used in the calculation of acoustic impedance.

**Step 4: Development of impedance map**

The impedance is calculated at sufficient points, sampled using Latin hypercube sampling, spread over the whole domain, and then interpolation through kriging is carried out to find the impedance over the complete domain, and an impedance map is generated to analyze the surface of shrimp scale.

## 5 Results and Discussion

### 5.1 Validation of the proposed framework through known material

The validity of the proposed algorithm has been rigorously tested and verified in the context of polyvinylidene fluoride (PVDF) as the primary material of interest, utilizing polyimide as the reference sample for comparison. The experimental process, detailed in Section 2, was meticulously described, recording the acoustic response of the system under study. In the experiment, the response has been obtained at multiple locations, however for brevity purposes, the results are presented for particular signals. The response signal is multifaceted, encompassing a spectrum of frequencies that evolve over time. To distill the essential characteristics of the signal, it becomes necessary to apply filtering techniques to identify the dominant frequencies within. The Short-Time Fourier Transform (STFT) and wavelet transform emerge as valuable tools for this purpose. The visualization of these frequencies after bandpass filtering, as extracted using STFT, is thoughtfully presented in Figure 4. This figure provides an insightful graphical representation of the frequency content of the signal, helping to elucidate and understand the dynamic changes in the system’s acoustic response over time. Further, the signal is decomposed through moDWT to distill the essential characteristics of the signal. While decomposing the signal into wavelet and scaling coefficients as described in Section 3.2, it is necessary to select the appropriate coefficients for the reconstruction of the filtered signal. Here, various time series are reconstructed from coefficients of different levels of decomposition, and then the time series which are having maximum energy content are selected for the complete reconstruction of the filtered signal. The time domain responses for both PVDF and Polyimide samples are illustrated in Figures 5(a and b), respectively. It provides a representation of the dynamic changes observed in the filtered responses due to the removal of unwanted frequencies from the signal through the maximal overlap discrete wavelet transform. The essential characteristics of the signal that are directly correlated to the properties of the specimen are extracted from this filtered response. However, the changes in the time domain are difficult to interpret and therefore the responses are transformed to the frequency domain where the predominant peak frequency is selected as a representative of the specimen characteristics as presented in Figure 6. Figure 6(a, b) illustrates the frequency spectra of the true responses of the reference sample PVDF and target medium polyimide. It serves as a foundation for impedance calculation, as it reveals the fundamental frequencies correlated to the material that helps us to deduce the reflectance property associated with a medium which ultimately provides the impedance of the target medium as described in Section 3.1.

**Figure 4.**
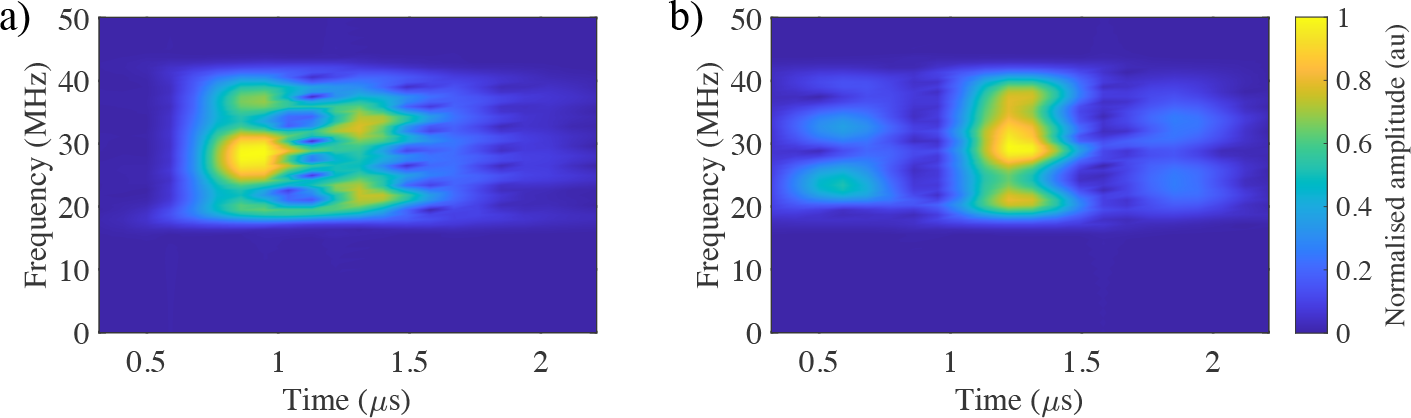
The figures display the Short-Time Fourier Transform (STFT), responses for (a) PVDF and (b) Polyimide, highlighting the presence of different frequencies in the signals that evolve over time, offering insights into their time-dependent characteristics.

**Figure 5.**
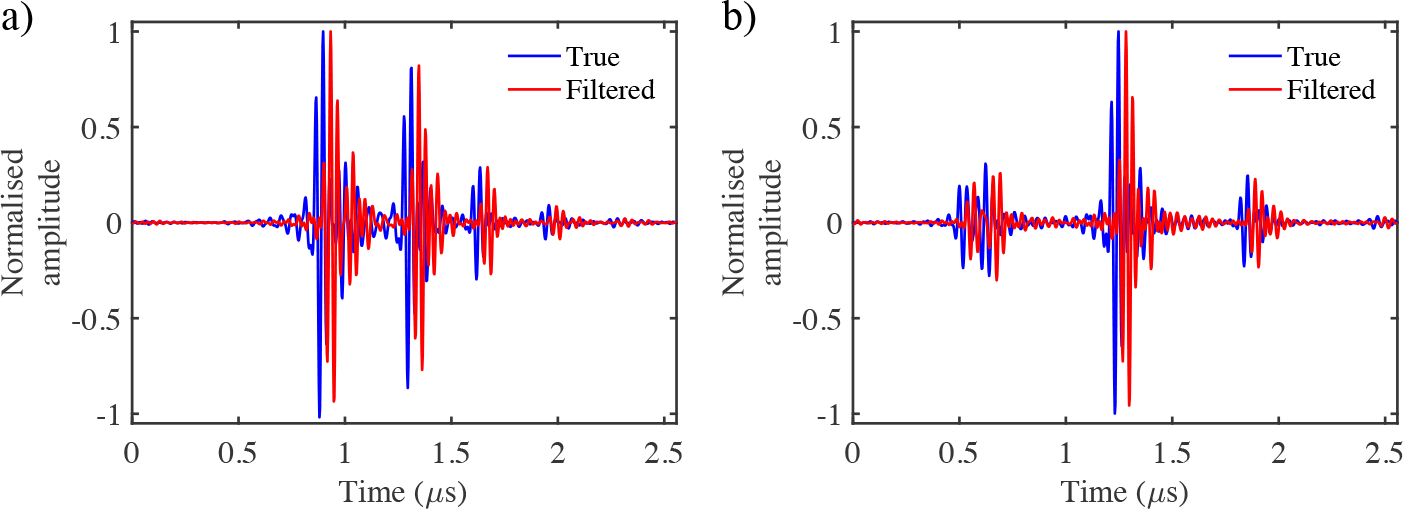
The figures represent the time domain responses for (a) PVDF and (b) Polyimide samples, demonstrate the dynamic changes in the filtered responses obtained through the modified discrete wavelet transform (moDWT), providing insights into the time-dependent behavior of the materials.

**Figure 6.**
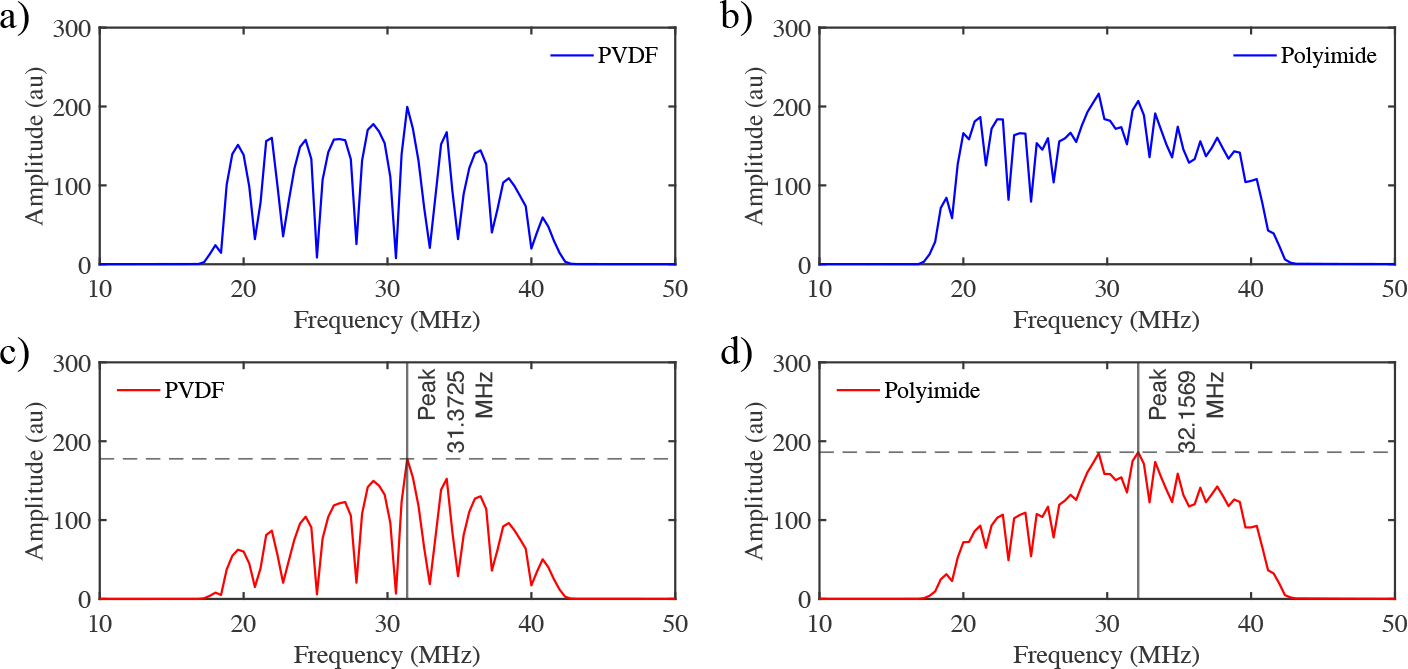
Displayed in the figures are the frequency spectra of the true responses (a) and (b), as well as the wavelet-transformed signals (c) and (d). These representations provide the identification of primary frequencies.

The accuracy of the complete algorithm is determined through accurate estimation of these frequencies. The frequency spectra of the filtered or wavelet transformed signal are shown in Figure 6 (c and d). The dominant peak from the frequency spectra of the filtered signal is obtained and shown in Figure 6 (c and d).

For the validation of the algorithm, let us consider the polyimide as the reference material, and the PVDF is our target material in this particular case, the substrate glass whose properties are known to us. After extraction of dominant frequencies, the impedance of the PVDF is calculated and the estimated value is compared with the true impedance value of the PVDF. The estimated impedance map generated of the PVDF through kriging using linear kernel function is shown in Figure 7 (a) and the relative error map is presented in Figure 7 (b). Table 1 shows the calculation of 10 spatial points selected using Latin hypercube sampling where the index in the table is anamously representing the test point’s location.

**Table 1.** Table 1 represents the corresponding frequency, measured and true impedance of the target and reference specimen and also their uncertainties.

|  |  |  |  |  |  |  |  |  |  |  |
| --- | --- | --- | --- | --- | --- | --- | --- | --- | --- | --- |
| Index(i) | 19 | 16 | 17 | 17 | 21 | 19 | 18 | 18 | 20 | 19 |
| Index(j) | 52 | 37 | 40 | 21 | 43 | 12 | 26 | 29 | 34 | 7 |
| $S_{target}$ (MHz) | 30.98 | 30.59 | 30.59 | 31.37 | 32.16 | 30.20 | 31.37 | 31.37 | 30.20 | 30.98 |
| $S_{ref}$ (MHz) | 32.16 | 31.77 | 31.77 | 32.55 | 32.94 | 32.16 | 32.16 | 32.94 | 32.16 | 32.94 |
| $S_{target}/S_{ref}$ | 0.96 | 0.96 | 0.96 | 0.96 | 0.98 | 0.94 | 0.98 | 0.95 | 0.94 | 0.94 |
| $Z_{sub}$ | 12.8 | 12.8 | 12.8 | 12.8 | 12.8 | 12.8 | 12.8 | 12.8 | 12.8 | 12.8 |
| $Z_{ref}$ | 3.31 | 3.31 | 3.31 | 3.31 | 3.31 | 3.31 | 3.31 | 3.31 | 3.31 | 3.31 |
| $Z_{pred}$ | 3.53 | 3.53 | 3.53 | 3.53 | 3.45 | 3.68 | 3.46 | 3.60 | 3.68 | 3.67 |
| $Z_{true}$ | 3.60 | 3.60 | 3.60 | 3.60 | 3.60 | 3.60 | 3.60 | 3.60 | 3.60 | 3.60 |
| Error % | 1.97 | 1.90 | 1.89 | 2.05 | 4.14 | 2.2 | 4.04 | 0.08 | 2.22 | 1.97 |

**Figure 7.**
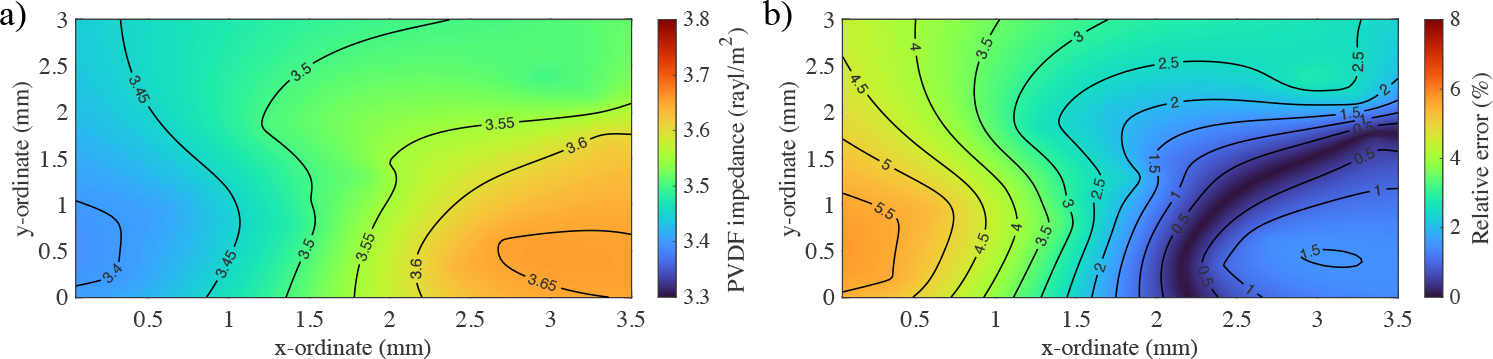
(a) Illustrates the distribution of estimated impedance values (PVDF or PI?), providing a comprehensive view of how impedance varies across the entire domain of interest. (b) demonstrates the relative error, which serves as a measure of the algorithm’s robustness and accuracy

It can be clearly shown that is error is within the 5% which demonstrates the efficacy and robustness of the proposed framework. The next section will discuss the application of the proposed framework in the domain of bio-cell which examines the impedance of the Shrimp scale in the present work.

### 5.2 Estimation and development of the impedance map of shrimp scale

In order to determine the acoustic impedance of the shrimp scale, we have applied the above-mentioned algorithm to analyze the acoustic response of the shrimp scale. For comparison and reference, we’ve used polyimide as the base material. The experimental methodology is already explained in detail in Section 2, where we meticulously conducted the experiment and recorded the acoustic response. The STFT highlights that dominant frequencies reflected through the shrimp scale are in the vicinity of 20 MHz to 40 MHz as shown in Figure 8. STFT helps in visualizing the time-frequency relation of the recorded acoustic response. This visualization serves as a tool for gaining a comprehensive understanding of how the acoustic characteristics of the shrimp scale change and develop over time, thereby contributing to insights into its dynamic behavior.

**Figure 8.**
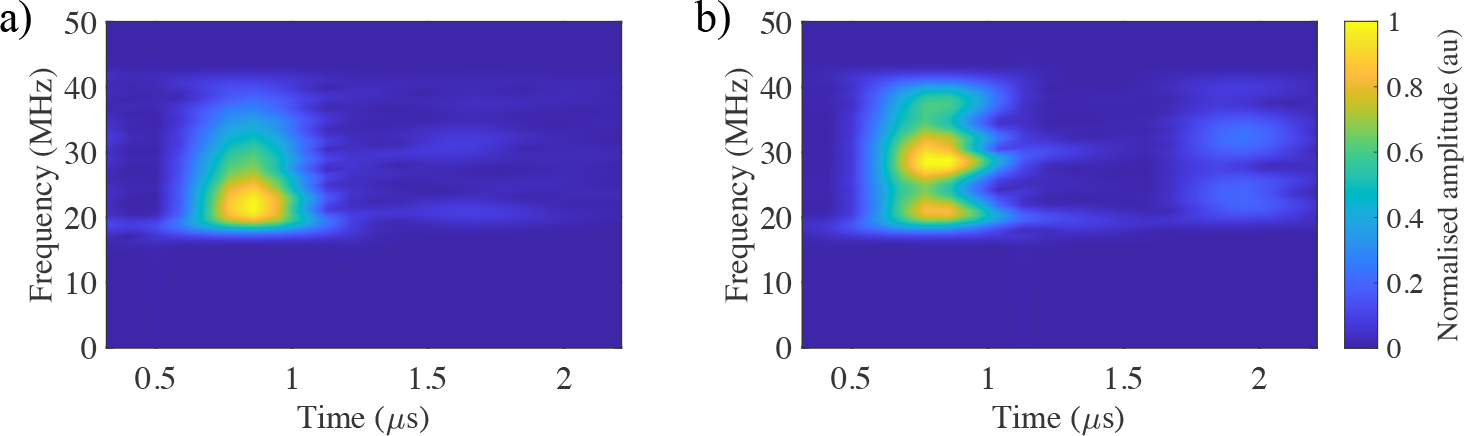
The figures display the Short-Time Fourier Transform (STFT), responses for (a) Shrimp scale and (b) Polyimide, highlighting the presence of different frequencies in the signals that evolve over time, offering insights into their time-dependent characteristics.

Figure 8 shows that there is a significant difference in the reflected frequency with time in both materials even though the transmitted signal is the same. This variation arises due to wave phenomenon primarily due to differences in the material impedance. The objective now is to isolate the dominant characteristics residing within it through moDWT. While decomposing the signal into wavelet and scaling coefficients as described in Section 3.2, the necessary coefficients of proper level having maximum energy are selected for the complete reconstruction of the filtered signal. Figure 9 (a and b) shows the time domain responses of the recorded signal and moDWT filtered signal of the Shrimp scale and polyimide. As already discussed in the previous case (PVDF and Polyimide). The essential characteristics of the signal that are directly correlated to the acoustic properties of shrimp scales can be extracted from this filtered response forming the fundamental basis for monitoring the bio-mechanical properties through signal processing.

**Figure 9.**
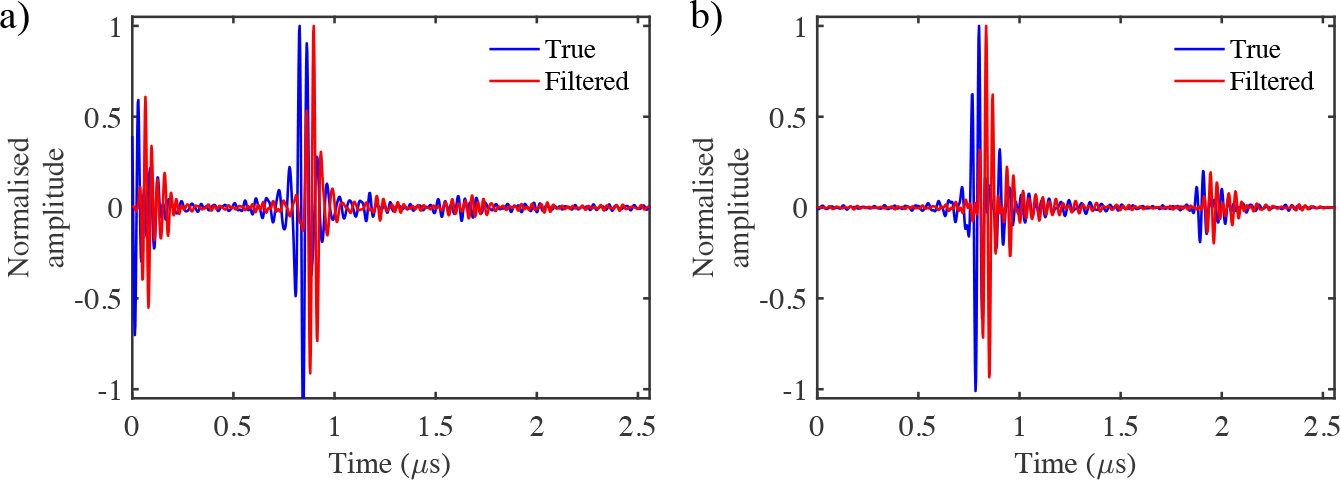
The figures represent the time domain responses for (a) Shrimp scale and (b) Polyimide samples, demonstrate the dynamic changes in the filtered responses obtained through the modified discrete wavelet transform (moDWT), providing insights into the time-dependent behavior of the materials.

However, the changes in the time domain are difficult to interpret and therefore the responses are transformed to the frequency domain where the predominant peak frequency is selected as a representative of the specimen characteristics as presented in Figure 10.

**Figure 10.**
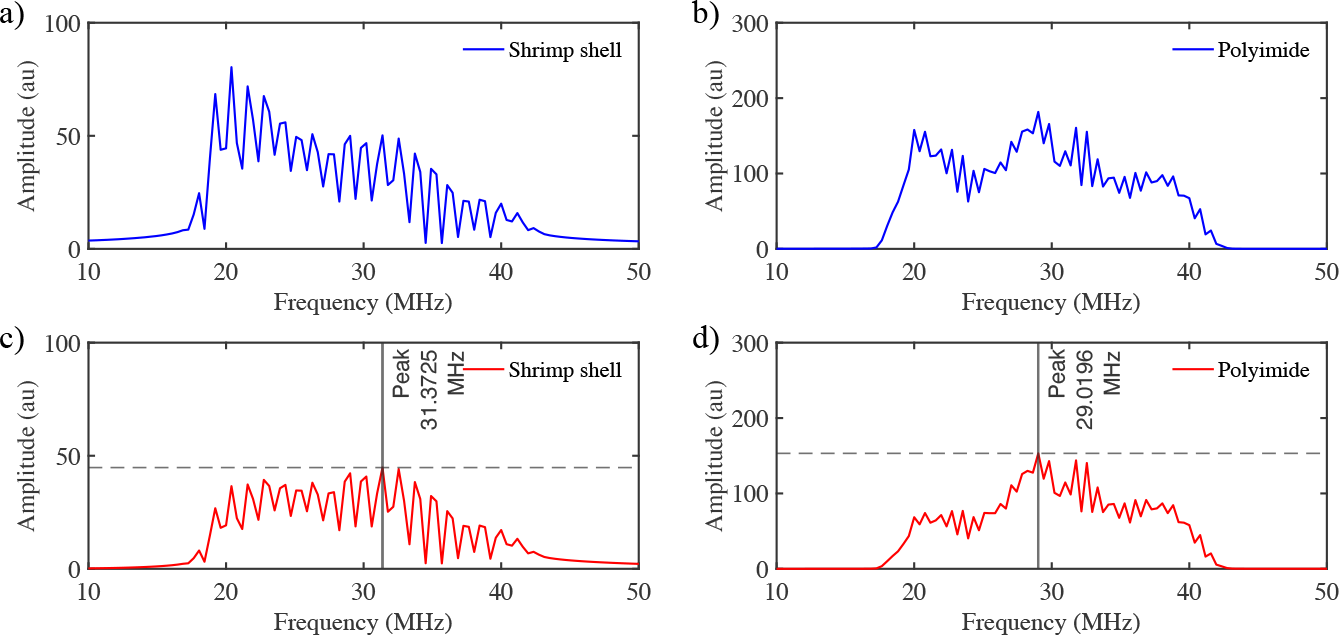
Displayed in the figures are the frequency spectra of the true responses (a) Shrimp scale or scale and (b) Polyimide, as well as the wavelet-transformed signals (c) and (d). These representations demonstrate the identification of primary frequencies which is a fundamental step in impedance calculation

Figure 10 (a and b) illustrates the frequency spectra of the true responses of the reference shrimp scale and target medium polyimide. The dominant frequency is selected which has the maximum amplitude in the spectra. It serves as a foundation for impedance calculation, as it reveals the fundamental frequencies correlated to the material that helps us to deduce the reflectance property associated with the medium which ultimately provides the impedance of the shrimp scale through the procedure described in Section 3.1. After extraction of dominant frequencies, the impedance of the shrimp scale is calculated and the impedance map is generated through kriging using the square exponential kernel function considering the greater variations in the Shrimp scale. The impedance map for the shrimp scale is shown in Figure 11 (a) and the standard deviation in the estimate is described in Figure 11 (b).

**Figure 11.**
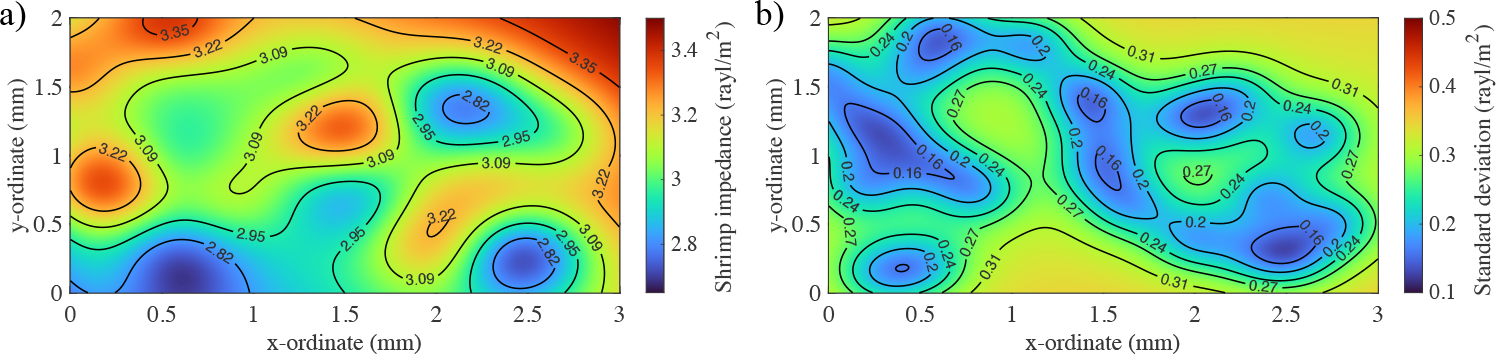
Figure (a)illustrates the distribution of estimated impedance values, providing a comprehensive view of how impedance varies across the entire domain of interest. Figure (b) demonstrates the standard deviation, which serves as a measure of uncertainty in the prediction

Figure 11 (a) represents the estimated impedance across the entire shrimp scale which offers a comprehensive insight. The figures visually represent the dispersion of estimated impedance values, providing a holistic understanding of how impedance fluctuates throughout the entire area of interest. This representation is indispensable for gaining insights into the impedance characteristics of the shrimp scale. Further, it can be inferred that the mean impedance values of the considered shrimp scale are between 2.8 to 3.4 rayl/*m*^2^ with an average standard deviation of 0.3. These values of impedance can now be used for any further investigations.

## 5 Conclusion

In this paper, a novel framework is proposed for estimating impedance through acoustic imaging applicable to various samples, including biological specimens like shrimp scales. The effectiveness of the framework is first established and confirmed through impedance characteristics of known materials. Here, PVDF serves as a reference point for calibration and ensuring the accuracy of our proposed algorithm. The results demonstrate the accuracy of the proposed algorithm is above 90%. Further, an impedance map is developed through the Gaussian process regression, which handles the complexity of the variations arising due to spatial variation in the biological specimen. The global analysis of acoustic impedance visualizations provides valuable insights into the functional aspects and complex biomechanical structure of various components within a shrimp’s exoskeleton. Overall, the presented innovative approach seeks to enhance our comprehension of shrimp biomechanics, potentially revolutionizing the detection of structural changes within marine communities.

## Acknowledgement

This work was supported by the Research Council of Norway, Cristin Project, ID: 2061348.

## Author contributions statement

A. H, A. S., and S. O. have conceptualized the idea. A. H. M. S., and K. A. designed and performed the experiments. S. O. implemented the algorithm with initial help from A. S. and A. H. Funding was secured by A. H. Formal analysis and experimental validation were performed by S.O. who also wrote the original draft, reviewed, and edited the manuscript with support from all co-authors.

## Data availability

The data sets used and/or analyzed during the current study are available from the corresponding author upon reasonable request.

